# Plasma proteomics in the UK Biobank reveals youthful brains and immune systems promote healthspan and longevity

**DOI:** 10.1101/2024.06.07.597771

**Authors:** Hamilton Se-Hwee Oh, Yann Le Guen, Nimrod Rappoport, Deniz Yagmur Urey, Jarod Rutledge, Anne Brunet, Michael D. Greicius, Tony Wyss-Coray

**Affiliations:** Graduate Program in Stem Cell and Regenerative Medicine, Stanford University, Stanford, CA, USA; The Phil and Penny Knight Initiative for Brain Resilience, Stanford University, Stanford, CA, USA; Wu Tsai Neurosciences Institute, Stanford University, Stanford, CA, USA; Department of Neurology and Neurological Sciences, Stanford University School of Medicine, Stanford, CA, USA; Quantitative Sciences Unit, Department of Medicine, Stanford University School of Medicine, Stanford, CA, USA; Department of Genetics, Stanford University, Stanford, CA, USA; Glenn Laboratories for the Biology of Aging, Stanford University, Stanford, CA, USA

## Abstract

Organ-derived plasma protein signatures derived from aptamer protein arrays track organ-specific aging, disease, and mortality in humans, but the robustness and clinical utility of these models and their biological underpinnings remain unknown. Here, we estimate biological age of 11 organs from 44,526 individuals in the UK Biobank using an antibody-based proteomics platform to model disease and mortality risk. Organ age estimates are associated with future onset of heart failure (heart age HR=1.83), chronic obstructive pulmonary disease (lung age HR=1.39), type II diabetes (kidney age HR=1.58), and Alzheimer’s disease (brain age HR=1.81) and sensitive to lifestyle factors such as smoking and exercise, hormone replacement therapy, or supplements. Remarkably, the accrual of aged organs progressively increases mortality risk while a youthful brain and immune system are uniquely associated with disease-free longevity. These findings support the use of plasma proteins for monitoring organ health and the efficacy of drugs targeting organ aging disease.

## MAIN

Aging leads to global dysfunction of organ systems, chronic disease, and ultimately death. Interventions including exercise, young plasma treatment, or rapamycin treatment seem to slow aging in model organisms, extending healthspan and lifespan^1^, but whether these findings would extrapolate to humans is unclear, as both our molecular understanding of human aging and our ability to efficiently assess the efficacy of aging interventions in humans is limited. Thus, molecular biomarkers of human biological age that are linked with age-related health and disease are needed.

Recent studies show that human organs age at different rates^2–5^ similar to what has been reported in animals^6–8^, which suggests a need for organ-specific measures of biological age. Previously developed organ age estimates include those developed from clinical metrics of organ function (glomerular filtration rate, blood pressure, etc), clinical blood chemistry, brain MRI scans, immune cell DNA methylation profiles, and the levels of organ-specific proteins in blood plasma^2–5^. Many questions regarding the reproducibility and utility of organ age estimates remain. For example, it is unclear the extent to which organ age estimates are stable across cohorts and longitudinal sampling, are sensitive to organ-specific diseases and modifiable lifestyle choices, and whether they predict mortality independent of each other and established aging biomarkers. Furthermore, it is unclear which organs are key to longevity in humans.

Given that plasma is simple to acquire, the levels of plasma proteins provide molecular information on organ function, and technologies to measure the levels of plasma proteins are rapidly advancing, we sought to further investigate the utility of plasma proteomics in estimating organ-specific biological age and understanding human longevity. In comparison to our previous study where we introduced our organ age estimation approach using SomaScan plasma proteomics data from 5,678 individuals^4^, here we aimed to test our approach in a much larger cohort of 44,526 individuals in the UK Biobank (age 40-70), using an orthogonal proteomics platform (Olink). The dramatically increased sample size and deeper tracking of age-related phenotypes allowed us to address several important questions in the field. Specifically, we assessed whether plasma protein-based age estimates for 11 major organs could identify different subtypes of organ agers in the population, predict risk of age-related organ diseases, track with beneficial/detrimental lifestyle choices, and predict mortality versus longevity (**Fig. 1a**).

**Figure 1.**
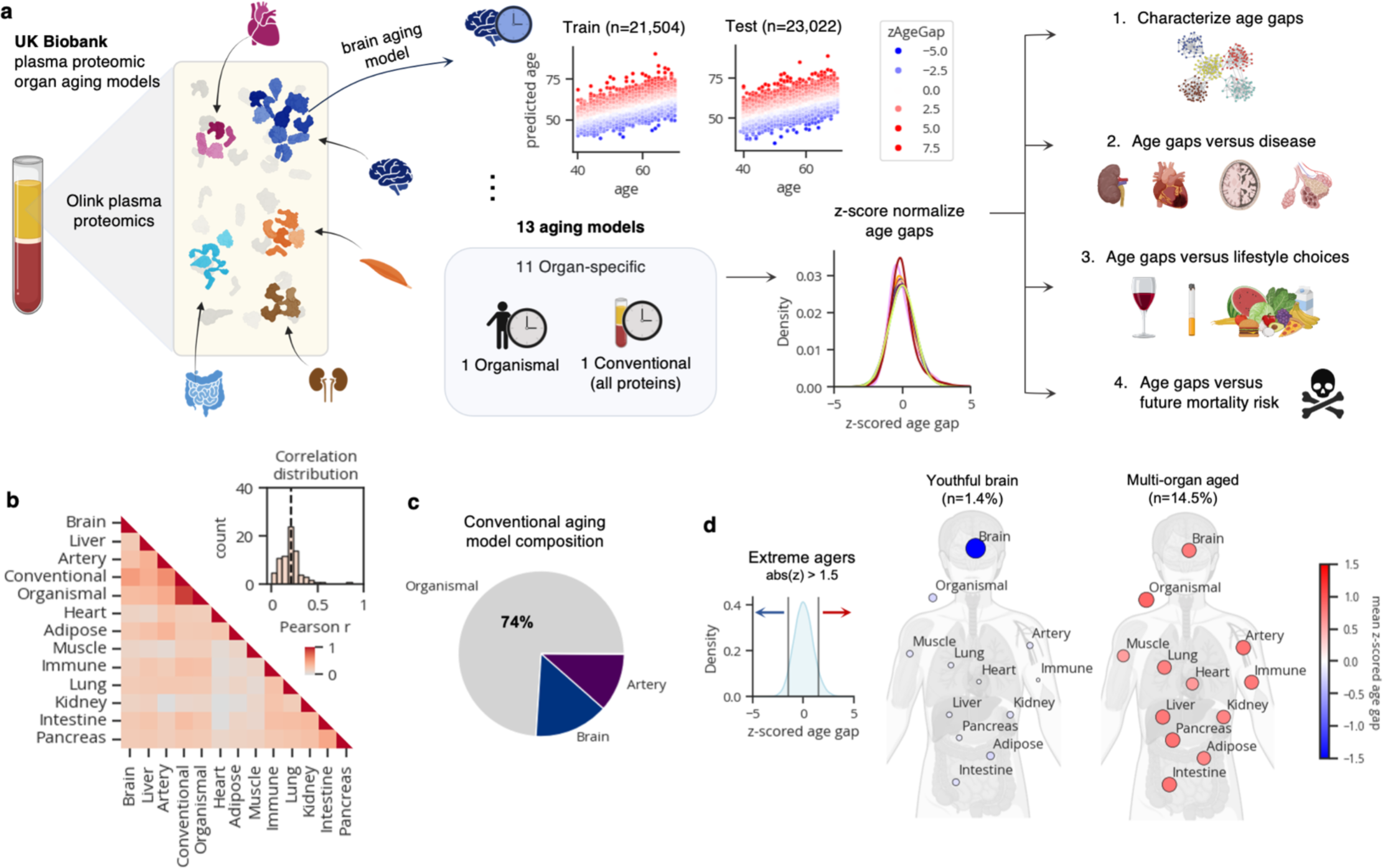
Plasma protein-derived organ age estimates in the UK Biobank. **a,** Study design to estimate organ-specific biological age from plasma proteomics data in the UK Biobank. A protein was called organ-specific if the gene encoding the protein was expressed at least four-fold higher in one organ compared to any other organ in the GTEx organ bulk RNA-seq atlas. Organ-specific protein sets were used to train LASSO chronological age predictors. Samples from 10/21 centers (n=21,504) were used for training and the remaining samples (n=23,022) were used for testing. An ‘organismal’ model, which was trained on the levels of non-organ-specific (organ-shared) proteins, and a ‘conventional’ model, which was trained on all proteins on the Olink assay, were also developed and assessed. Model age gaps were calculated and then z-score normalized per organ to allow for direct comparisons across organs. Age gaps were characterized (Fig. 1), and tested for associations with disease risk (Fig. 2), modifiable lifestyle choices (Fig. 3), and mortality risk (Fig. 4). **b**, Pairwise correlation of organ age gaps from all samples. Inset histogram shows the distribution of all pairwise correlations, with the dotted line representing the mean. **c,** A Lasso regression model was used to predict conventional age based on organ ages and organismal age. Organismal, brain, and artery ages were sufficient to predict conventional age with r^2^=0.97. Relative weights are shown as a pie chart. **d**, Extreme agers were defined by a 1.5 standard deviation increase or decrease in at least one age gap. The mean organ age gaps of extremely youthful brain agers and accelerated multi-organ agers are shown.

### Plasma protein-derived organ age estimates in the UK Biobank

To derive measures of organ-specific physiological state and biological age from the plasma proteome, we 1) identified plasma proteins likely derived from a specific organ (**Supplementary Tables 1-2**), 2) trained a machine learning model to predict chronological age based on the levels of identified organ-specific proteins (**Supplementary Tables 3-4**), and 3) calculated the age gap based on each person’s predicted age (the relative predicted age compared to individuals of the same chronological age; See methods; **Fig. 1a**). The age gap provides a measure of relative biological age compared to same-aged peers.

We performed these three steps for each of 11 major organs including adipose tissue, artery, brain, heart, immune tissue, intestine, kidney, liver, lung, muscle, and pancreas. To compare organ age gaps to organ-agnostic measures of biological age, we also derived age gaps from an “organismal” aging model trained on non-organ-specific proteins and a “conventional” aging model trained on all proteins on the Olink assay. We confirmed the top proteins in the conventional aging model overlapped with a previous proteomic aging model developed on the UK Biobank dataset^9^.

Data from ten of twenty-one plasma collection centers were used for model training and the remaining eleven for testing (**Fig. 1a**). Model performance was highly stable across train and test centers (**Extended Data Fig. 1a-b**). Age gaps were z-scored per aging model to allow for direct comparison between organs in downstream analyses (**Fig. 1a**). We observed some sex differences between organ age gaps with males having older kidneys, immune systems, and intestines, while females had older adipose tissue, arteries, and hearts (**Extended Data Fig. 1c-d**).

After deriving organ age gaps, we first determined the uniqueness of each measure. If organs truly aged at different rates, then the age gap of one organ should be independent from the age gap of another in the same individual. Thus, we calculated pairwise correlations between organ age gaps. As expected, based on previous literature, we found organ age gaps were only mildly correlated (mean r=0.21; **Fig. 1b**). Of note, organ age gaps were largely different from the conventional age gap, suggesting they capture information not measured by conventional proteomic aging models in the field. Instead, the conventional age gap was strongly correlated with the organismal age gap (r=0.87). Estimated organismal, brain, and artery age were sufficient to explain 97% of the variance in estimated conventional age, with organismal age carrying 74% of the weight (**Fig. 1c**).

We previously identified extreme organ agers who displayed especially fast aging in just a single organ or many organs^4^. We searched for extreme agers in the UK Biobank by identifying individuals with age gaps above or below 1.5 standard deviations from the population average in any organ (top and bottom ∼6-7% percentiles). Indeed, we found groups of fast (>=1.5 z-age gap) and slow (<= –1.5 z-age gap) organ agers who had extreme age gaps in only a single organ (1-2% of samples each, 33% total; **Fig. 1d**; **Extended Data Fig. 1e**). We also identified multi-organ agers who had two or more extreme organ age gaps (26% of samples**; Extended Data Fig. 1e**). 14% of samples were ambiguous, with both positive and negative extreme age gaps and 27% of remaining samples were defined as normal agers (**Extended Data Fig. 1e**). Multi-organ agers were significantly older than normal and single organ agers, suggesting aged organs accumulate over time (**Extended Data Fig. 1f-g**). Youthful agers were not identified in our previous study^4^ based on SomaScan data, potentially due to limited sample size and detection of different proteins between platforms.

We next sought to determine the longitudinal stability of organ age gaps over several years within an individual. In other words, does a person with a biologically older heart at age 50 still have an older heart at age 60? We analyzed a subset of 937 individuals who had plasma proteomics data from two to three visits, spanning 1 to 15 years from baseline. We assessed the correlations between age gaps at baseline versus visit two, for each organ. We found medium to strong correlations across visits (mean r=0.6), suggesting age gaps are relatively stable, while also likely reflecting dynamic alterations to biological age based on lifestyle and disease as well as technical variability (**Extended Data Fig. 2a**).

Examining longitudinal stability of extreme ager status, we found that individuals who were extreme agers in a given organ at visit 1 were 3-22 times more likely to be an extreme ager in that same organ at visit 2, compared to individuals who were not extreme agers in that organ at visit 1 (**Extended Data Fig. 2b**). Still, a considerable proportion (68%) of visit 1 extreme agers were no longer categorized as extreme agers for their respective organs in visit 2. Therefore, we examined changes in age gap status by bins separated by 0.5 standard deviations up to +/-1.5, rather than binary extreme ager status. We found that the vast majority of visit 1 extreme agers stayed within the same signed bins across visit 2 (76%) and visit 3 (72%), suggesting that the overall directionality of their age gaps was stable (**Extended Data Fig. 2c-e**).

Together, these data show that organ-specific biological age estimates can be derived from plasma proteomic data in the UK Biobank, and that each organ age estimate provides unique information about the individual, perhaps related to organ-specific health. Moreover, within 2 independent visits, deviation from chronological age tends to be stable within individuals.

### Organ age estimates predict future age-related disease

For an estimate of biological age to be informative, it must robustly associate with the physiological state of the organ or individual and consequently, with age-related health and disease outcomes. Hence, we sought to determine whether organ age gaps could predict future diseases in their respective organs. We tested the associations between all 13 z-scored age gaps and 15 incident age-related diseases (2-17 year follow-up) using Cox proportional hazard regression, while adjusting for age and sex. Following Benjamini-Hochberg correction for multiple hypothesis testing, we identified 176 positive and 4 negative significant associations out of 195 tests (**Fig. 2a; Supplementary Table 5**).

**Figure 2.**
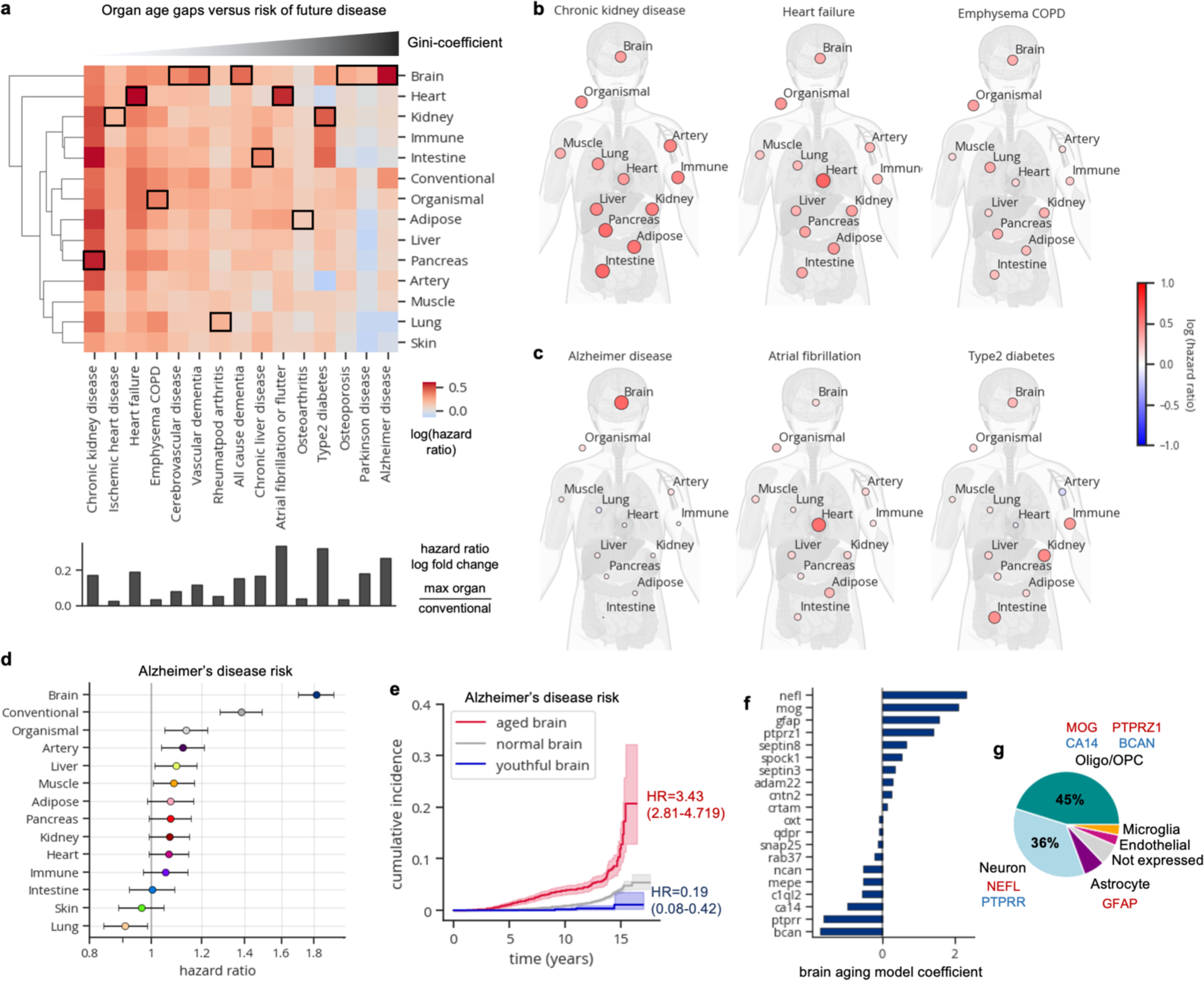
Organ age estimates predicts future age-related disease. **a,** Cox proportional hazards regression was used to test the association between age gaps and future disease risk, adjusted for age-at-blood-draw and sex. Heatmap colored by age gap log(hazard ratio) is shown. Heatmap columns are ordered by the Gini-coefficient of age gaps per disease. The most significant associations per disease are highlighted with black borders. The conventional age gap was never the most significant. The log fold change in hazard ratios between the organ with the most significant age gap versus the conventional age gap is shown below the heatmap. **b,** Body plots showing log hazard ratios from the heatmap in **a,** are shown for diseases of systemic aging. **c,** Body plots showing log hazard ratios from the heatmap in **a,** are shown for diseases of single-couple organ aging. **d,** Forest plot visualizing the results from the heatmap in **a,** for Alzheimer’s disease risk. Age gap hazard ratios and 95% confidence interval shown. **e,** Cumulative incidence plot showing increased risk of Alzheimer’s disease in extreme accelerated brain agers and decreased risk in youthful brain agers. **f,** Bar plot displaying the top 10 protein coefficients in the brain aging model. **g,** Pie chart displaying proportion of brain aging proteins assigned to each brain cell-type based on single-cell RNA-sequencing. Cell type was assigned based on cell type with the maximum expression of a given gene.

We discovered highly significant associations between heart aging and atrial fibrillation (hazard ratio [HR]=1.75, q<1x10^–230^) and heart failure (HR=1.83, q=1.39x10^–230^), pancreas aging (HR=1.77, q=5.03x10^–220^) and kidney aging (HR = 1.64, q = 5.25x10^–217^) with chronic kidney disease, brain aging with Alzheimer’s disease (HR=1.81, q=2.29x10^–72^) and lung aging with chronic obstructive pulmonary disease (HR=1.39, q=7.20x10^–32^). Liver aging was associated with chronic liver disease (HR=1.18, q=4.31x10^–11^), albeit the strength of the association was modest and similar compared to other organs. Importantly, organ-specific age gaps consistently exhibited stronger predictive power than conventional age gaps across all diseases (**Fig. 2a**).

The widespread significant associations between organ aging (176/195) and disease underscore the systemic nature of aging. While certain diseases may have notably systemic causes, others may stem primarily from dysfunction in a single organ. Identifying the degree of systemic versus local contributions in disease onset could yield insights into biological mechanisms and prevention strategies. Thus, for each disease, we calculated the Gini-coefficient, a measure of statistical dispersion (originally intended to measure income inequality), of organ age gap log hazard ratios.

Interestingly we found that chronic kidney disease, ischemic heart disease, heart failure, and chronic obstructive pulmonary disease (COPD) were predicted by aging of many organs, suggesting systemic underpinnings (**Fig. 2b**). Whether these diseases are caused by systemic aging or the prodromal phases of these diseases induce aging across the body is unclear. Conversely, Alzheimer’s disease was predicted primarily by brain aging (**Fig. 2c-d**). Other more organ-specific diseases included atrial fibrillation which was primarily predicted by heart aging and type 2 diabetes which was predicted primarily by kidney (HR=1.58, q=2.54x10^–206^), intestine (HR=1.54, q=1.18x10^–126^), and immune (HR=1.49, q=1.03x10^–103^) aging (**Fig. 2c**).

We also investigated the associations between extreme organ age and disease risk (**Extended Data Fig. 3a; Supplementary Table 6**). Individuals with more than one aged organ had significantly increased risk for nearly every disease we examined. Individuals with only aged brains, hearts, kidneys, or lungs also showed broad increased risk of disease. On the other hand, individuals with 2-4 youthful organs were protected from some disease including chronic kidney disease, osteoarthritis, and COPD; however, individuals with 5-7 youthful organs were not globally significantly protected and instead had increased risk for diabetes and Parkinson’s disease, suggesting a youthful appearing aging signature is not always beneficial. Among youthful organ profiles, only the brain and immune system were protective for at least three diseases (nominal p-value<0.05) while not conferring increased risk of any other diseases. A youthful brain was especially protective of Alzheimer’s disease (HR=0.13, p=0.046, q=0.15) and other dementias, while a youthful immune system was protective of diabetes (HR=0.52, p=0.0073, q=0.044), atrial fibrillation, and cerebrovascular disease (**Extended Data Fig. 3a**).

We further examined all individuals with youthful or aged brains – regardless of single or multi-organ ager status – and their associations with Alzheimer’s disease risk (**Fig. 2e**). We found that individuals with aged brains had a 3.4-times increased risk (HR=3.43, p=7.26x10^–34^) while those with youthful brains had a striking 81% reduced risk (HR=0.19, p=4.21x10^–5^) of Alzheimer’s disease compared to those with normal aging brains (**Fig. 2e**). Regarding sample size, 121 of 2,618 individuals (4.6%) with aged brains developed Alzheimer’s disease over 17 years, while only 6 of 2,002 individuals (0.3%) with youthful brains developed the disease.

The specificity of the association between brain aging and dementia led us to determine whether organ age gaps were associated with brain volume based on magnetic resonance imaging (MRI) data from follow-up visits (**Extended Data Fig. 3b**). We confirmed that brain aging was uniquely associated with increased volume of the ventricles and decreased volume of cortical regions.

We also assessed organ age gap associations with disease progression, by regressing age gaps against years since diagnosis, for individuals who were diagnosed with disease before blood draw (**Extended Data Fig. 3c**). We found that many organ age gaps increased throughout chronic kidney disease progression. Interestingly, the brain age gap was not associated with dementia progression (**Extended Data Fig. 3d**), suggesting the brain aging model uniquely captures age-related changes leading up to dementia but not after.

Given these robust associations, we sought to gain further insights into organ aging by examining aging model proteins and their weights (**Extended Data Fig. 4a**). The strongest weighted protein in the brain aging model was neurofilament light chain (NEFL; **Fig. 2f**), which increases with age and is a clinical biomarker of neurodegeneration that is often measured in clinical trials for Alzheimer’s disease^10,11^ and was recently approved as a surrogate endpoint for a clinical trial to treat superoxide dismutase 1 amyotrophic lateral sclerosis (SOD1-ALS)^12,13^. Our data suggest it may also be a viable surrogate endpoint for functional deterioration of the brain, risk of dementia, and brain aging. Other highly weighted brain aging proteins include myelin oligodendrocyte protein (MOG), a component of the outer surface of myelin sheaths, and glial fibrillary acidic protein (GFAP) a marker of reactive astrocytes, which both increased with age, as well as brevican (BCAN), a brain extracellular matrix component produced by oligodendrocyte precursor cells, and protein tyrosine phosphatase receptor type R (PTPRR), which decreased with age (**Fig. 2f**). Plasma NEFL, GFAP, and BCAN were previously highlighted as predictors of future dementia risk^14^. Model weights between UK Biobank-Olink and SomaScan^4^ brain aging models were moderately correlated (Spearman ρ=0.5), with NEFL, BCAN, and PTPRR being most aligned in effect and directionality across platforms (**Extended Data Fig. 5a**). Using the permutation feature importance for biological aging (FIBA) algorithm^4^, we found that all of the top seven most highly weighted brain aging proteins contributed to the prediction of Alzheimer’s disease risk, showing the importance of leveraging the information from many brain-derived proteins to understand brain aging and disease (**Extended Data Fig. 5b**).

We then sought to determine which cell types these proteins were likely derived from by analyzing public human brain single-cell RNA-sequencing data^15^ (**Extended Data Fig. 5c-d**). Interestingly, we found that around half of brain aging proteins were specific to the oligodendrocyte lineage, with the rest expressed mostly in neurons, then astrocytes (**Extended Data Fig. 5d**), pointing to white matter as a key aging region, as suggested by human brain MRI and mouse brain RNA-sequencing studies^16,17^.

Lung aging was explained primarily by lysosome-associated membrane glycoprotein 3 (LAMP3), a protein expressed specifically in type II alveolar stem cells, secretoglobin family 1A member 1 (SCGB1A1) also known as club cell secretory protein (CCSP), a marker of club cells, and C-C Motif Chemokine Ligand 18 (CCL18), a cytokine expressed by alveolar macrophages, potentially reflective of stem cell dysfunction and inflammation in the lung with age. Heart aging was explained primarily by N-terminal pro b-type natriuretic peptide (NT-proBNP), a vasodilating hormone that increases in response to heart damage, while kidney aging was explained by renin (REN), a protein involved in blood pressure regulation. Both NT-proBNP and REN were previously identified as key heart and kidney aging proteins, respectively, based on SomaScan plasma proteomics data^4^ and are well-established biomarkers of heart and kidney function. All aging model protein weights are provided in **Supplementary Table 4** and the top 20 proteins for each model are shown in **Extended Data Fig. 4a**.

Together, these data show that plasma protein-derived organ age estimates are linked with age-related organ diseases and can reveal insights into the aging biology of their respective organs.

### Organ age estimates are sensitive to modifiable lifestyle choices

We next explored whether biological age estimates grounded in physiological states of organ function are sensitive to changes in lifestyle. We tested the associations between all 13 z-scored age gaps and 15 lifestyle factors (diet, alcohol, smoking, exercise), adjusted for age and sex using linear regression (**Supplementary Table 7**). We found 66 positive and 79 negative significant associations after correcting for multiple hypothesis testing. In line with their known health impacts, smoking, alcohol, and processed meat intake were associated with age acceleration across several organs, while “vigorous exercise” and oily fish consumption were associated with youthful organs (**Fig. 4a**).

**Figure 3.**
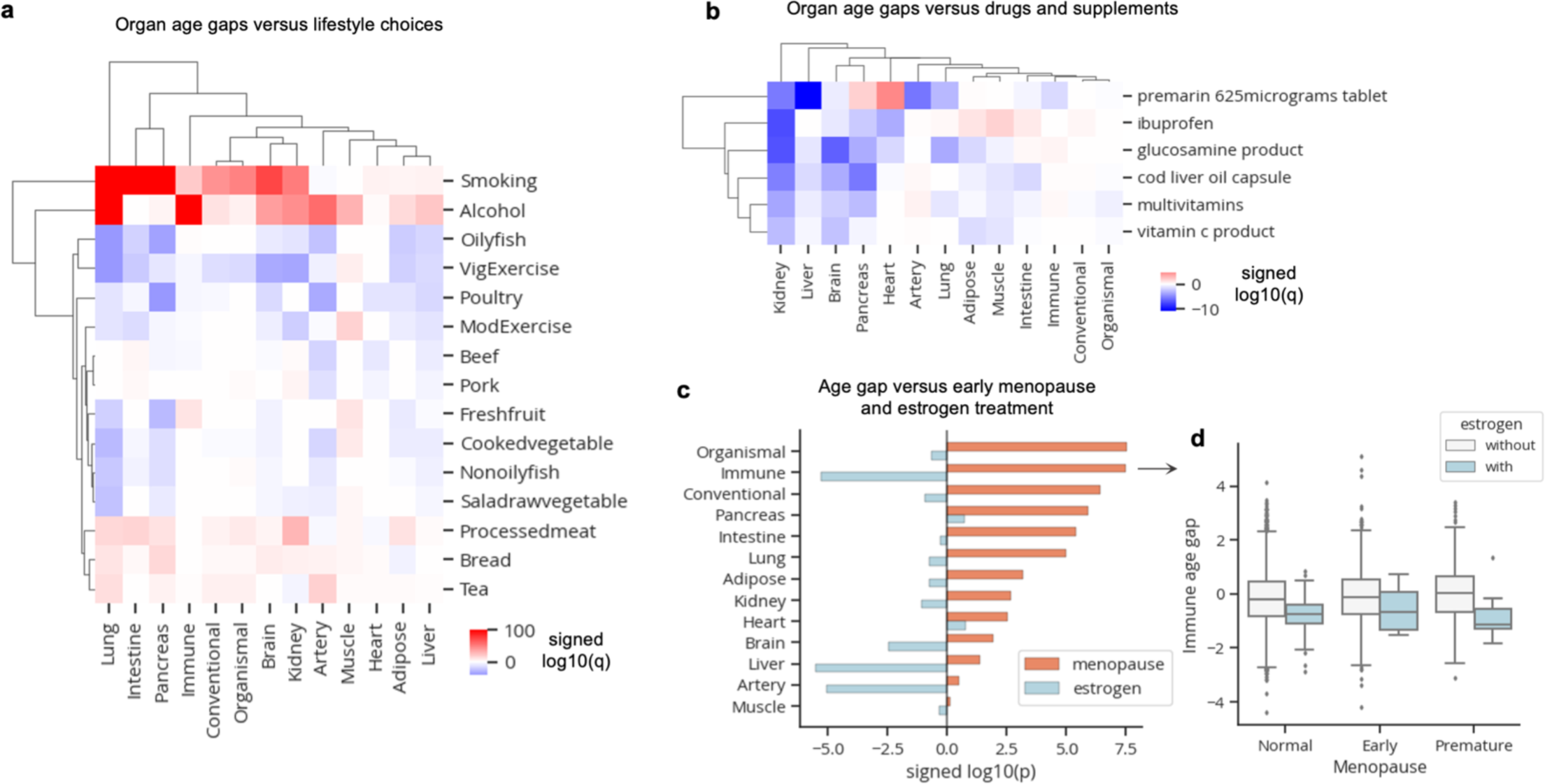
Organ age estimates are sensitive to modifiable lifestyle choices. **a**, Linear regression was used to determine the association between age gaps and modifiable lifestyle choices while accounting for age and sex. Heatmap colored by signed log10(q-value) is shown. Only significant (q<0.05) values are colored. **b,** Linear regression was used to determine the association between age gaps and drugs/supplement intake while accounting for age and sex. Only drugs with significant (q<0.05) youthful associations in at least two organs are shown. Heatmap is colored by signed log10(q-value). Only significant (q<0.05) values are colored. **c,** Linear regression was used to determine the association between age gaps versus early menopause and estrogen treatment together. Bar plot showing signed log10(p-value) for menopause and estrogen covariates is shown. **d,** Boxplot visualization of immune age gaps in individuals stratified by menopause status and estrogen treatment. Standard boxplot structure was used.

**Figure 4.**
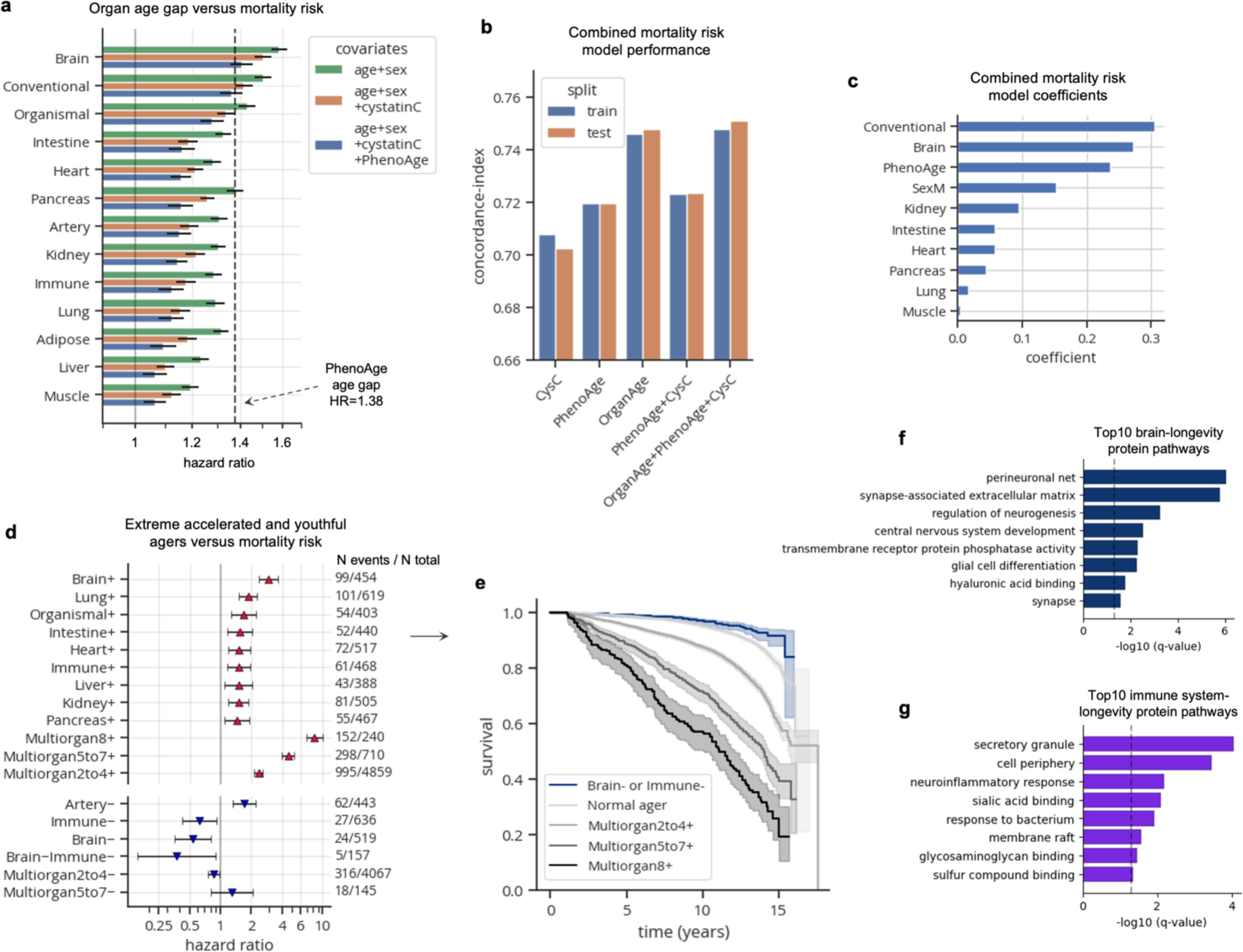
Accrual of aged organs progressively increases mortality risk while brain and immune system youth is associated with longevity. **a**, Bar plot showing results from Cox proportional hazards regression analyses, testing the associations between age gaps and future all-cause mortality risk, controlling for age, sex, (and blood cystatin C; and PhenoAge). Hazard ratios and 95% confidence intervals are shown. PhenoAge age gap hazard ratio (1.38) is shown as a dotted line for reference. **b,** Concordance indices from various Lasso-regularized Cox proportional hazard models trained to predict mortality risk. Performance across train and test centers is shown. Covariates for each model, in addition to age-at-blood-draw and sex, are labeled on the x-axis. **c,** Model coefficients shown for the combined model (OrganAge+PhenoAge+CysC) from **b**. **d,** Forest plot showing results from Cox proportional hazards regression, testing the associations between extreme ager status and future all-cause mortality risk, controlling for age-at-blood-draw and sex. Only significant (q<0.05) associations are shown. Age gap hazard ratios, 95% confidence intervals, number of events out of the total sample size are shown. **e,** Kaplan-Meier curves showing survival over 17-year follow-up of normal agers, progressive levels of multi-organ agers (2-4, 5-7, 8+ aged organs), and individuals with youthful brains or immune systems. **f-g,** Gene ontology pathway enrichment results (with all genes as background) from top ten brain (**f**) and immune (**g**) aging proteins by mortality risk FIBA score.

We also tested the associations between all 13 z-scored age gaps and consumption of 137 drugs/supplements (n cases>=100; **Supplementary Table 8**). After multiple hypothesis test correction, we found 5 products – Premarin, ibuprofen, glucosamine, cod liver oil, multivitamins, vitamin C – that were significantly (q<0.01) associated with more youthful age in at least two organs (**Fig. 4b**). Ibuprofen, glucosamine, cod liver oil, multivitamins, and vitamin C products were associated with youth primarily in the kidneys, brain, and pancreas (**Fig. 4b**).

Premarin is a conjugated estrogen medication typically prescribed to women experiencing post-menopausal symptoms, and estrogen medication has been recently shown to be associated with reduced mortality risk in the UK Biobank^18^. Thus, we wondered whether estrogen medications may extend longevity by preventing menopause-induced accelerated aging of organs. We grouped all post-menopausal estradiol/oestrogen medications together and identified 47 women with normal, early, or premature menopause (but not late menopause) who were treated with estrogen by the time of blood draw. We subsetted our analysis to women among these menopausal groups and tested the independent associations of age at menopause and estrogen treatment with organ age gaps using linear regression. Interestingly, women with earlier menopause were age accelerated across nearly all organs, in line with the well-documented adverse health consequences of early menopause^19^ (**Fig. 4c**). Conversely, estrogen treatment was associated with more youthful immune systems, livers, and arteries suggesting intervention with estrogen treatment may protect these organs from menopause-induced age acceleration resulting in extended survival (**Fig. 4d**).

Though these cross-sectional findings should be interpreted with caution, they underscore the sensitivity of plasma protein-derived organ age estimates to various lifestyle choices with known health impacts and their potential utility in assessing the effects of novel interventions in more rigorous experimental settings.

### Accrual of aged organs progressively increases mortality risk while brain and immune system youth is associated with longevity

We next sought to determine whether organ age estimates were associated with future mortality risk. We tested associations between organ age gaps and all-cause mortality risk, adjusting for age and sex, over a 2-17 year follow-up using Cox proportional hazards regression (**Supplementary Table 9**). We found significant associations in all organs with a standard deviation increase in age gaps conferring 20%-60% increased risk of death (**Fig. 4a**), similar to findings from SomaScan-based organ aging models^4^. These associations were robust to additional adjustment with blood cystatin-C, a marker of kidney filtration rate, and PhenoAge, an established blood biochemistry/cell counts-based biological age estimate (whose age gap has a mortality risk hazard ratio of 1.38 in the UK Biobank), suggesting that organ age estimates provide independent information not captured by existing clinical biomarkers. Surprisingly, brain aging was most strongly predictive of mortality (HR=1.58, p=4.40x10^–291^) (**Fig. 4a**), suggesting that the brain may be a central regulator of lifespan in humans similar to findings in animal models (worms, flies, mice)^20–22^. Indeed, individuals with aged brains had increased risk for several diseases beyond dementia including COPD and heart failure (**Extended Data Fig. 3a**) consistent with previous studies showing that the brain regulates systemic inflammation^23–27^.

These data suggested that organ age estimates may provide additional predictive power for mortality, beyond cystatin-C and PhenoAge. We compared the concordance-index, a metric of Cox model performance equivalent to area under the curve (AUC), of regularized Cox proportional hazard models trained on cystatin-C alone, PhenoAge alone, organ ages alone, and combinations of these aging biomarkers, with chronological age and sex as covariates in each model. We found that the model with all variables had the strongest performance, comparable to the performance of the model with organ ages alone (**Fig. 4b**), suggesting that organ ages not only provide additional predictive power, but also that they sufficiently capture the information from other clinical aging biomarkers. Examination of the combined model coefficients revealed that conventional age, brain age, PhenoAge, and biological sex were the strongest independent predictors of mortality (Fig. 4c). Application of FIBA to understand contributions of brain and conventional aging model proteins on mortality risk highlighted BCAN, NEFL, and PTPRR from the brain as well as ectodysplasin A2 receptor (EDA2R, organismal protein), chemokine C-X-C motif ligand 17 (CXCL17, organismal protein), and elastin (ELN, artery protein) from the conventional aging model as important proteins (**Extended Data Fig. 6a-d**).

While each organ age gap was associated with risk of death, we wondered whether the accrual of aged organs would be increasingly detrimental. Hence, we tested whether single-organ extreme agers and multi-organ agers (bins of 2-4, 5-7, 8+ organs) had an increased risk of death compared to normal agers, while adjusting for age and sex (**Supplementary Table 10**). Interestingly, we found that while having a single aged organ (brain, lung, intestine, heart, immune, kidney, liver, pancreas) conferred a 1.5-3 times increased risk, having 2-4, 5-7, and 8+ extremely aged organs conferred a 2.3-, 4.5-, and 8.2-times (q=3.52x10^–70^, q=1.11x10^–104^, and q=1.72x10^–123^) increased risk of death, respectively (**Fig. 4d-e**). Notably, over 60% of individuals with 8+ extremely aged organs at blood draw died within 15 years (**Fig. 4d**).

We then sought to determine whether youthful organ profiles were associated with longevity (**Supplementary Table 10**). We found individuals with youthful appearing arteries had increased mortality risk, and those with multi-organ youth had no difference in mortality risk compared to normal agers (**Fig. 4d**). Using FIBA, we found that artery-protein thrombospondin 2 (THBS2), a protein that decreases with age but has a positive association with mortality risk, was responsible for the non-linear association between the artery age gap and mortality risk (**Extended Data Fig. 6c**; **Extended Data Fig. 4a**). Why individuals with broad multi-organ youth are not protected is unclear.

Notably though, individuals with youthful brains (HR=0.53, q=6.58x10^–3^) and immune systems (HR=0.63, q=0.034) had significantly reduced mortality risk (**Fig. 4d**), similar to their unique protective associations with disease (**Extended Data Fig. 3a**). We therefore assessed individuals with both youthful brains and immune systems and found that this group was most strongly protected from mortality (HR=0.37, p=0.027; **Fig. 4d-e**), underscoring the additive longevity benefits of having a resilient brain and immune system. Regarding sample size, 792 of 10,000 individuals (7.92%) of normal agers died within 17 years, while only 5 of 157 individuals (3.2%) with youthful brains and immune systems died.

To probe the underlying mechanisms implicated in brain- and immune system-related longevity, we performed gene ontology enrichment analyses of the top ten brain and immune aging model proteins based on mortality risk FIBA scores (**Extended Data Fig. 6f-g**; **Fig. 4f-g**). Selection of brain- or immune system-specific plasma proteins as background for the enrichment test did not result in significant associations, so we used all genes as background. Perineuronal net was the most enriched brain pathway (BCAN, PTPRZ1, NCAN) and secretory granule was the most enriched immune system pathway. Intriguingly, neuroinflammation was another enriched immune system pathway, composed of metalloproteinase-9 (MMP9), a regulator of perineuronal net composition, as well as tumor necrosis factor receptor superfamily member 1B (TNFSRF1B), and integrin alpha M (ITGAM) also known as CD11B. Youthful brain agers had higher levels of BCAN and NCAN and youthful immune agers had lower levels of MMP9 (**Extended Data Fig. 6f-g**), suggesting that global preservation of brain extracellular matrix partly due to reduced degradation by peripheral inflammatory factors and reduced chronic inflammation partly regulated by the brain could be crucial for promoting longevity.

Together, these data reveal that plasma-protein derived organ age estimates improve upon existing aging biomarkers for mortality risk prediction, the accrual of aged organs progressively increases mortality risk, and a youthful brain and immune system are key to longevity.

## DISCUSSION

Our findings based on plasma proteomics data (∼3,000 proteins) from nearly 45,000 individuals in the UK Biobank establish plasma protein-derived organ age estimates as robust indicators of organ age, health, and disease/mortality risk beyond gold-standard clinical aging biomarkers and reveal key proteins implicated in the aging process. Furthermore, we show that organ age estimates are stable across train/test centers and longitudinal visits and are cross-sectionally associated with modifiable lifestyle choices and therapeutic use, and thus, lay the foundation for human experiments testing the effects of novel longevity interventions on the biological age of organs at the individual level.

Though we define correlates of organ aging that explain heterogeneity in disease and mortality, future work is needed to understand the molecular and environmental determinants and sequence of organ aging. Our discovery of multi-organ agers and their increased prevalence with age (**Fig. 1d, Extended Data Fig. 1g**) suggests that aged organs likely accumulate over time within an individual, but how and why is unclear. Consistent longitudinal sampling combined with comprehensive patient phenotyping regarding environmental stresses and genetic background are needed to elucidate the dominant sequences and causes of organ aging at the population and individual levels. Improved technologies that allow for the sampling of thousands of more molecules (proteins, lipids, and metabolites) from human tissue will aid in obtaining a more global understanding of human aging heterogeneity and its causes.

Furthermore, additional studies are needed to understand why individuals with youthful profiles in many organs are not protected from disease and mortality (**Fig. 4d, Extended Data Fig. 3a**). While this may be commonly assumed to be true, numerous studies have shown non-linear associations between clinical biomarkers of health (ie. BMI, blood ALT, platelet count) and mortality risk with extreme “youthful” phenotypes being associated with higher risk^28–30^. This observation emphasizes the complexities of biological age estimation and the importance of investigating non-linear associations between aging signatures with disease and mortality to derive interpretable and actionable insights.

Here, we find evidence that the brain and immune system may be central regulators of aging and longevity in humans, as accelerated brain aging is most strongly predictive of earlier mortality and a youthful brain and immune system is most predictive of longevity (**Fig. 4a-d**). After all, the brain regulates numerous critical age-related functions throughout the body including circadian rhythm, blood pressure, energy homeostasis, and stress response via the neuroendocrine and autonomic nervous systems, and chronic inflammation has been heavily implicated in aging^1,31^. Intriguingly, recent studies show bidirectional communication between the brain and immune system in aging and disease relevant contexts, such as chronic stress, atherosclerosis, and infection^23–27^. These observations suggest that accelerated aging or maintenance of youth in the brain and immune system likely has broad age-related effects across the body, though additional studies are needed to test this hypothesis more rigorously in humans.

Regarding the molecular alterations that occur with brain aging, we find many unexpected age- and disease-associated changes in oligodendrocyte lineage- and extracellular matrix-proteins, implying extensive changes beyond neuroinflammation and neurodegeneration. Indeed, myelin degeneration and defective remyelination with mouse aging causes cognitive deficits^32^ and aggravates Alzheimer’s disease pathology^33^, and APOE4, the strongest genetic risk factor for late-onset Alzheimer’s disease, impairs myelination via cholesterol dysregulation in oligodendrocytes^34^. Moreover, the observation that white matter regions exhibit the most pronounced shifts in aging microglial transcriptomes^17^, coupled with the enrichment of genetic risk variants for neurodegenerative diseases in microglial and oligodendrocyte genes^35^, underscores a potential link between oligodendrocyte aging and neuroinflammation, and their relevance to neurodegeneration. Future studies exploring these multi-cellular interactions in the aging brain and their interactions with the periphery may reveal key insights into human health and longevity.

## METHODS

### Participants

The UK Biobank is a population-based prospective cohort with ‘omics and phenotypic data collected on approximately 500,000 participants, aged 40 to 69 years at recruitment between 2006 and 2010. A subset of these had subsequent visits after the initial assessment 20,337 participants had a first repeat assessment (2012-2013), approximately 85,000 had a first MRI imaging visit (from 2014 to ongoing), and approximately 9,000 had a second MRI imaging visit (from 2019 to ongoing). Details on available phenotypes can be found online at https://biobank.ndph.ox.ac.uk/showcase/ and all participants provided informed consent. A subset of these participants had their blood sample processed with the Olink proteomics assessed, which selection, data processing and quality control are described in (Sun et al., 2023, https://www.nature.com/articles/s41586-023-06592-6). Briefly, 53,014 samples were from the initial assessment visit, 1,172 were from the first imaging visit, and 1,123 were from the second imaging visit.

We defined participants’ last-known-age as either age-at-death, or the difference between the latest date available in ICD9, ICD10, operating procedure, cancer registry or UKB assessment visit fields and birth date, this age corresponds to censoring in the following survival analyses. Additionally, we define the age-at-onset of several groups of diseases based on a combination of information in ICD10, ICD9, operating procedure, and cancer registry. The following disease groups were defined, as in (https://www.medrxiv.org/content/10.1101/2023.09.13.23295486v1.full), Type-2 diabetes (E11), Ischemic heart disease (I20 to I25), Cerebrovascular disease (I60 to I69), Chronic liver disease (K70, K73 to K76), Chronic kidney disease (N18), All cause dementia (A81, F00 to F03, F05, F10, G30, G31, I67), Vascular dementia (F01, I67), Alzheimer’s disease (F00, G30), Parkinson’s disease and parkinsonism (G20 to G22), Rheumatoid arthritis (M05, M06), Macular degeneration (H35), Osteoporosis (M80, M81), Osteoarthritis (M15 to M19), Prevalent hypertension (I10 to I13, I15), Colorectal cancer (C18, C19, C20), Lung cancer (C33, C34), Esophageal cancer (C15), Liver cancer (C22), Pancreatic cancer (C25), Brain cancer (C71), Leukemia (C91, to C95), Non-Hodgkin lymphoma (C82 to C86), Breast cancer (C50), Ovarian cancer (C56, C57), Prostate cancer (C61). For heart-related diseases the group definitions were based on https://www.medrxiv.org/content/10.1101/2023.12.19.23300218v1, Heart failure (ICD9: 4254, 4280, 4281, 4289, ICD10: I110, I130, I132, I255, I420, I428, I429, I500, I501, I509), and Atrial fibrillation or flutter (ICD9: 4273, ICD10: I480, I481, I482, I483, I484, I489, OPCS4: K571, K621, K622, K623, K624, X501, X502).

Medications reported in the verbal interview (https://biobank.ctsu.ox.ac.uk/crystal/label.cgi?id=100075) were also analyzed with a minimum of 100 participants per medication.

Brain MRI-derived phenotypes extracted with Freesurfer (https://biobank.ctsu.ox.ac.uk/crystal/label.cgi?id=110) were analyzed, and age-at-MRI as well as estimated-total-intracranial volume were regressed out.

UK Biobank data were analyzed under Application Number 45420.

### Identification of organ-specific plasma proteins

We used the same methods we developed in our previous study^4^ to identify organ-specific plasma proteins. Briefly, we identified organ-enriched genes: genes that were expressed at least four times higher in a single organ compared to any other organ based on human tissue bulk RNA-seq data from the Gene Tissue Expression Atlas (GTEx). We refer to our previous study for details. Our classification of organ-enriched genes are provided in **Supplementary Table 1**. We mapped these genes to proteins measured by Olink to identify organ-specific plasma proteins in the UK Biobank dataset (**Supplementary Table 2**).

### Organ age estimation

We used LASSO regression models to build chronological age predictors, a.k.a. aging models, to estimate biological age. The LassoCV function from the scikit-learn^36^ Python package was used to identify the optimal lambda parameter value using 5-fold cross validation. The lambda value that achieved 95% of the performance of the highest performing lambda value was applied to scikit-learn’s Lasso function to derive sparse aging models. Each organ aging model was trained using a distinct set of organ-specific plasma proteins. An organismal aging model was trained using non-organ-specific proteins. A conventional aging model was trained using all proteins. Age gaps were calculated as the residual of predicted age linearly regressed against actual age. Age gaps were z-scored per aging model to normalize for differences in age prediction accuracy. Z-scored age gaps were used for all analyses. Extreme agers were defined as individuals with an age gap z-score > 1.5 or z-score < –1.5 in a given aging model. Conventional age gaps were not included in the extreme ager analyses due to their high similarity to organismal age gaps (**Fig. 1b-c**).

Aging models were trained and tested on 44,526 baseline assessment plasma samples which had measurements for 2,923 proteins from Olink Explore proteomics. 7 proteins with missing values in over 10% of samples were removed, leaving 2,916 proteins for model development. Remaining missing values were imputed using scikit-learn’s KNNImputer function. The knn-imputer was trained on 21,504 samples from 10 randomly selected sample collection centers (train centers: 11013, 11009, 11014, 11008, 11018, 11007, 11017, 11005, 11002, 11023) and tested on 23,022 samples from the remaining 11 centers (test centers: 11010, 11011, 11016, 11020, 11021, 11004, 11003, 11006, 11012, 11001, 11022, 10003). The number of neighbors used for the knn-imputer was set to the square root of the sample size of the training dataset (n=147). Protein values were z-score normalized based on the means and standard deviations of proteins in the training split (**Supplementary Table 3**) prior to model training. Aging models were trained and tested using the same train-test split as the knn-imputer. Aging models performed near equally across train and test samples (**Extended Data Fig. 1b**; **Fig. 4b**), demonstrating robustness of missing value imputation and biological age estimation. Aging model weights are provided in **Supplementary Table 4**.

### Longitudinal age gap analyses

Longitudinal age gap analyses (**Extended Data Fig. 2**) required use of plasma proteomics data collected across multiple visits from the same individual. However, proteomics data from post-baseline samples were obtained from an earlier version of the Olink assay with 1,459 proteins and were not compatible with models trained on the ∼3k protein platform. Therefore, for longitudinal age gap analyses, we trained a distinct set of organ aging models using the subset of ∼1.5k proteins that were measured across all visits.

1,463 proteins were measured across all visits. 4 proteins with missing values in over 10% of samples were removed, leaving 1,459 proteins for model development. Missing values for baseline samples were knn-imputed as described above. Post-baseline samples were not imputed to prevent biased imputation towards baseline data from the same individual. 1.5k-protein-based aging models were trained on 44,406 baseline samples from individuals who did not have follow-up proteomics data. Importantly, samples from individuals with longitudinal proteomics data were not included in model training to prevent model training-evaluation contamination. 1k-protein-based aging models were LASSO regression models trained to predict the predicted age from the 3k protein-based organ aging models. The lambda value that achieved 90% of the performance of the highest performing lambda value was used for sparsity. Recursive feature elimination using scikit-learn’s RFECV function was additionally used to further simplify the models to maximize the number of testable samples with unimputed data. Liver and muscle 1k-based aging models were removed due to low correlation (r<0.8) with 3k-based aging models. Models were tested and evaluated on longitudinal data from 1,176 unique individuals who had non-missing values for all remaining aging model proteins (880 baseline, 843 imaging-visit-1, and 786 imaging-visit-2 samples). The mean number of years between imaging-visit-1 and baseline was 9.1 years (st. dev.=1.8), and the mean number of years between imaging-visit-2 and imaging-visit-1 was 3.3 years (st. dev.1.6).

### Statistical analyses

Cox proportional hazards regression (CoxPHFitter function from lifelines^37^ Python package) was used to assess the associations between organ age gaps and future disease or mortality risk. Linear regression (OLS function from statsmodels^38^ Python package) was used to assess the associations between organ age gaps and lifestyle factors recorded at the time of blood draw. All Cox and linear regression models included age and sex as additional covariates. Multiple hypothesis testing correction was applied, when appropriate, using the Benjamini-Hochberg method, and the significance threshold was a 5% false discovery rate. Corrected p-values are referred in the manuscript as q-values. Results and sample sizes for these statistical tests are provided in **Supplementary Tables 5-10**.

## DATA AVAILABILITY

Organ age estimates for all UK Biobank participants will be returned to the UK Biobank and available through Showcase.

## CODE AVAILABILITY

Organ aging models can be accessed in a Python package called organageUKB (at the time of publication) to easily estimate organ age from any Olink plasma proteomics sample. All aging model weights are provided in **Supplementary Table 4**. Means and standard deviations to z-score protein levels before applying model weights are provided in **Supplementary Table 3**.

## AUTHOR CONTRIBUTIONS

H.S.O., J.R., and T.W.-C conceptualized the study. H.S.O. led study design and analyses. Y.L.G. led data processing of UK Biobank clinical phenotypes and advised on analyses. N.R. aided in longitudinal organ age gap analyses and figure generation. D.Y.U. aided in disease progression analyses and figure generation. J.R., A.B., and M.D.G. provided key insights. H.S.O. produced figures and wrote the manuscript. T.W.-C. edited the manuscript. T.W.-C supervised the study. All authors critically revised the manuscript for intellectual content. All authors read and approved the final version of the manuscript.

## ACKNOWLEDGEMENTS

We thank B. Lehallier, A. Antebi, L. Gold, and members of the Wyss-Coray laboratory for feedback and support and D. Channappa for laboratory management. This work was supported by the Stanford Alzheimer’s Disease Research Center (National Institute on Aging grants P50AG047366 and P30AG066515), the National Institute on Aging (AG072255, T.W.-C), the Milky Way Research Foundation (T.W.-C.), and the Knight Initiative for Brain Resilience (T.W.-C.). It is further supported by the Stanford Graduate Fellowship (H.S.O.), the National Science Foundation Graduate Research Fellowship (H.S.O.), and the NIH Center for Clinical and Translational Education and Research award, Biostatistics, Epidemiology and Research Design (BERD) Program (UL1TR003142, Y.L.G.).

---

**Extended Data Figure 1.**
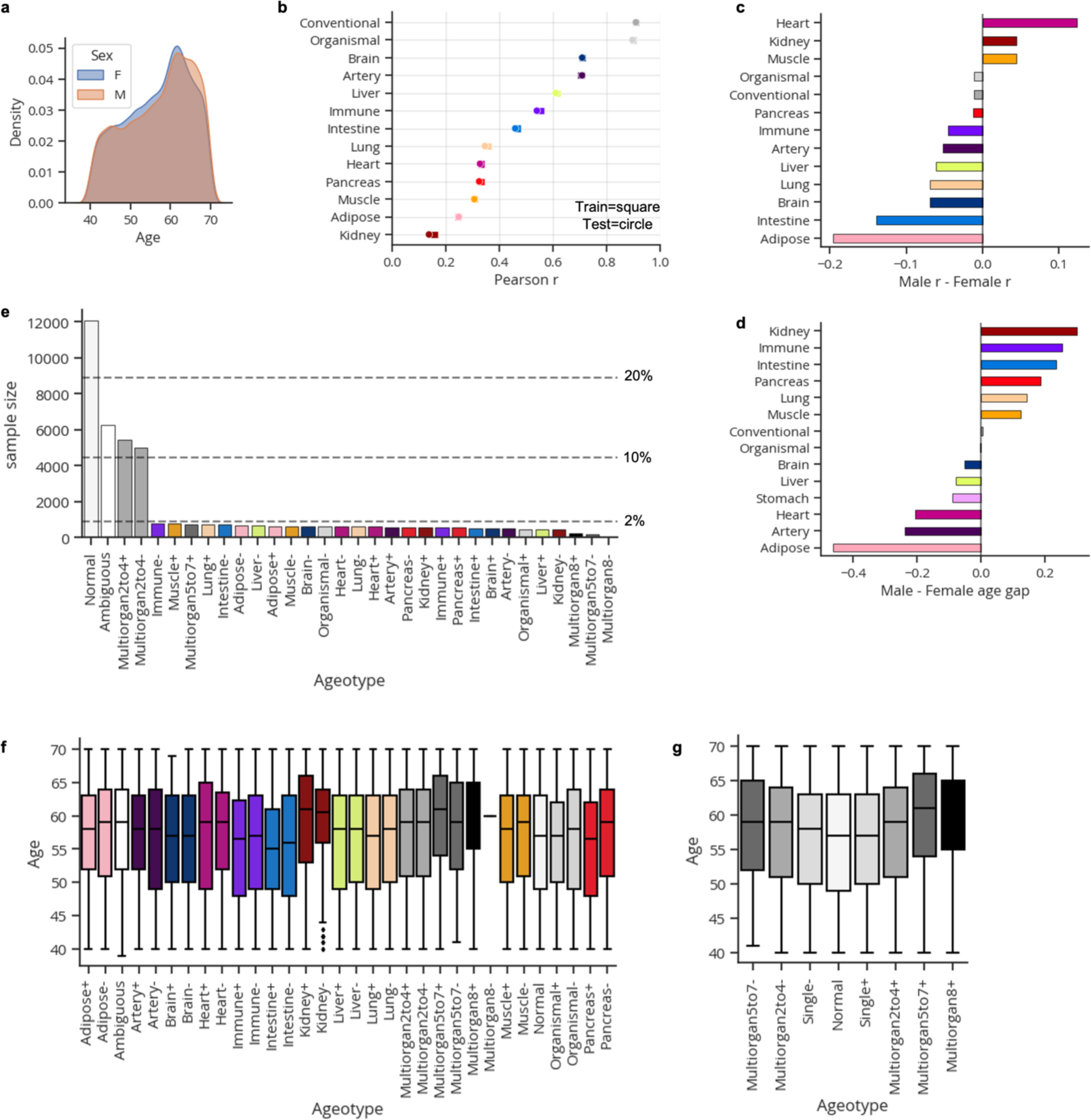
Organ aging models in the UK Biobank. **a,** Age-at-blood-draw distribution by biological sex. **b,** Correlation between predicted and actual age across all aging models and train/test splits. **c,** Difference in correlation between predicted and actual age by biological sex. **d,** Mean difference in organ age gaps between males and females. **e,** Extreme ager ageotype sample sizes and proportions. **f,** Age distributions per extreme ager ageotype **g,** Age distributions per aggregated extreme ager ageotype. Individuals with many aged or youthful organs are significantly older than normal and single organ agers.

**Extended Data Figure 2.**
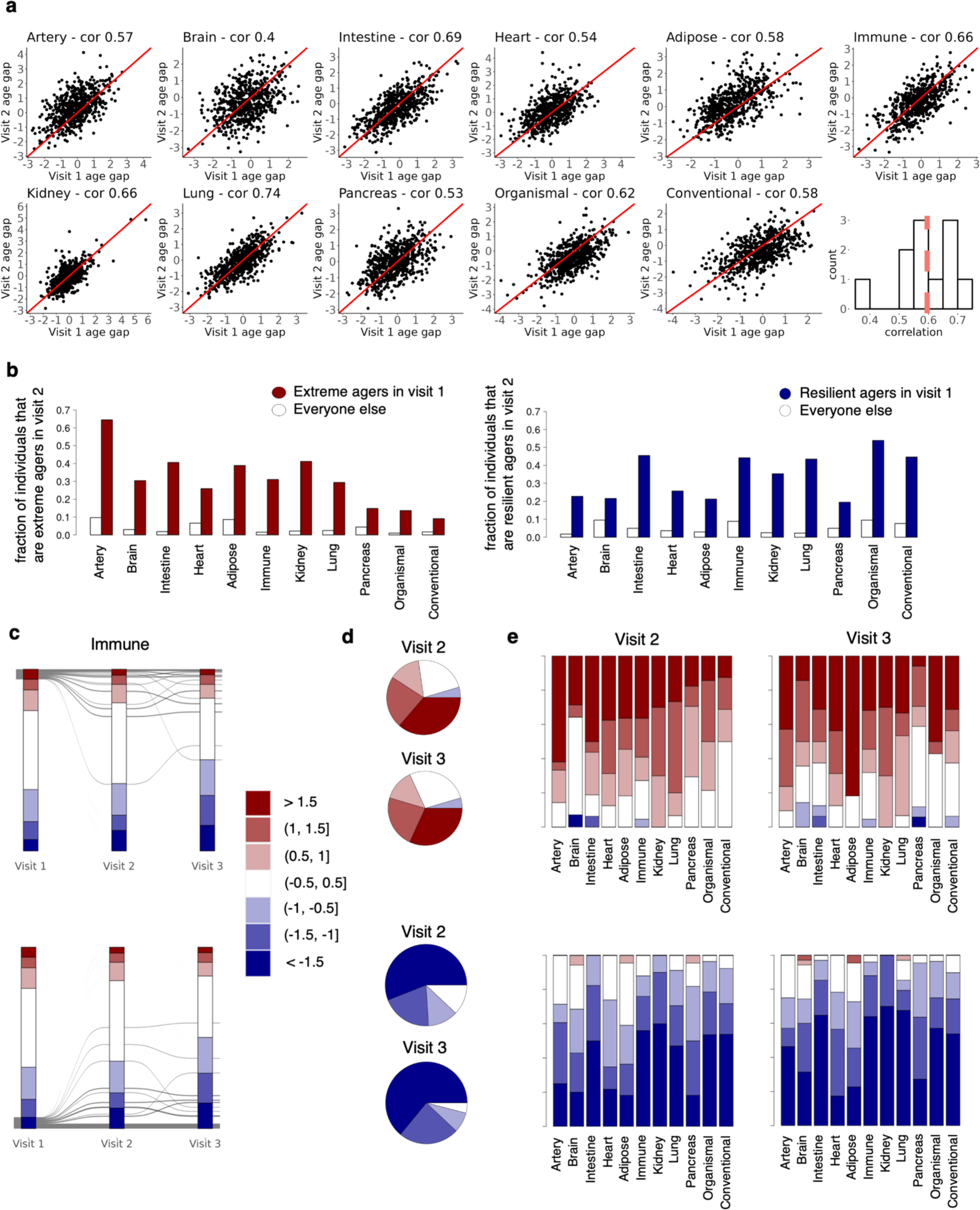
Age gaps are stable across longitudinal visits. **a,** Longitudinal proteomics data from a subset of 937 individuals were analyzed. Longitudinal data were available only on the 1k-protein platform, so new aging models trained on the 1k-platform were developed. New aging models were trained on 44,406 samples without longitudinal data and tested on samples with longitudinal data (937 unique individuals). Only 1k-aging models with age estimates that were correlated r^2^>=0.8 with 3k-based age estimates were included for downstream analyses. Correlation between visit 1 (baseline, 2006-2010) and visit 2 (imaging visit 1, 2014+) age gaps are shown. **b,** Bar plot showing fractions of visit 1 extreme agers and non-visit 1 extreme agers that are extreme agers in the same organ in visit 2. Equivalent plot for youthful agers is shown on the right. **c,** Age gaps were grouped into bins of 0.5 standard deviation to determine changes in age gap bins across visits. Individual trajectories across visits for extreme immune agers are shown. Equivalent plot for youthful immune agers is shown at the bottom. **d,** Pie chart showing percent distribution of immune age gap bins in visit 2 (2014+) and visit 3 (2019+) for individuals who are extreme immune agers in visit 1. Equivalent plot for youthful immune agers is shown at the bottom. **e,** Stacked bar plot showing percent distribution of age gap bins in visit 2 and visit 3 for individuals who are extreme agers in visit 1. Equivalent plot for youthful agers is shown at the bottom.

**Extended Data Figure 3.**
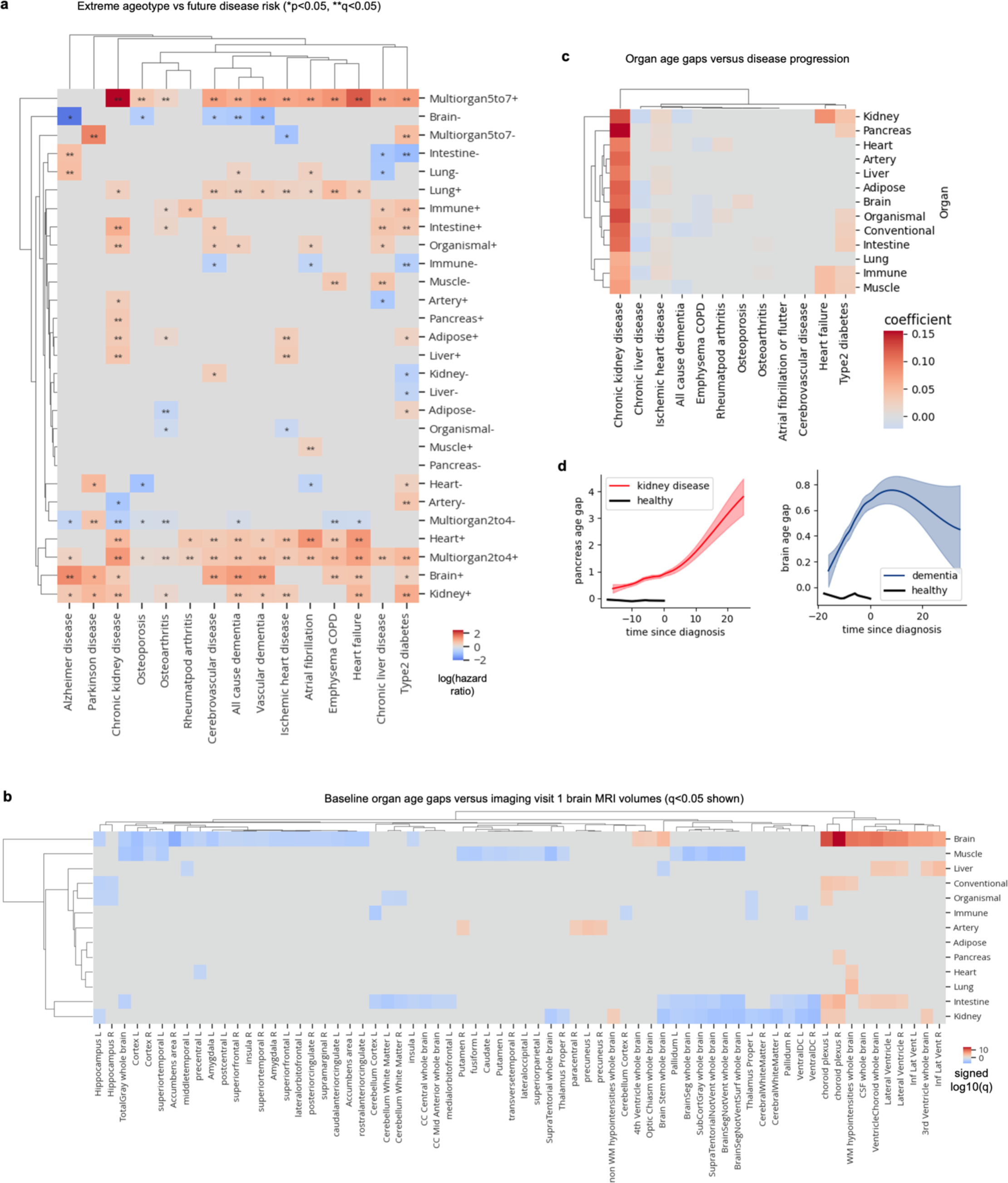
Ageotypes versus disease risk and age gaps versus brain volume. **a,** Cox proportional hazards regression was used to determine the association between extreme ageotypes and future disease risk, controlling for age and sex. Heatmap colored by age gap log(hazard ratio) is shown. *p<0.05, **q<0.05. Non-significant hazard ratios (p<0.05) were set to zero. **b,** Linear regression was used to determine the association between baseline organ age gaps and imaging visit 1 brain MRI volumes, controlling for age-at-blood-draw, age-at-imaging-visit-1, sex, and estimated total intracranial volume. Non-significant effect sizes (q<0.05) were set to zero. Red indicates positive associations, while blue indicates negative associations. **c,** Linear regression was used to determine the association between organ age gaps and years since disease diagnosis. Non-significant effects (q<0.05) were set to zero. **d,** Visualization of results from **c**. Organ age gap versus years since diagnosis shown.

**Extended Data Figure 4.**
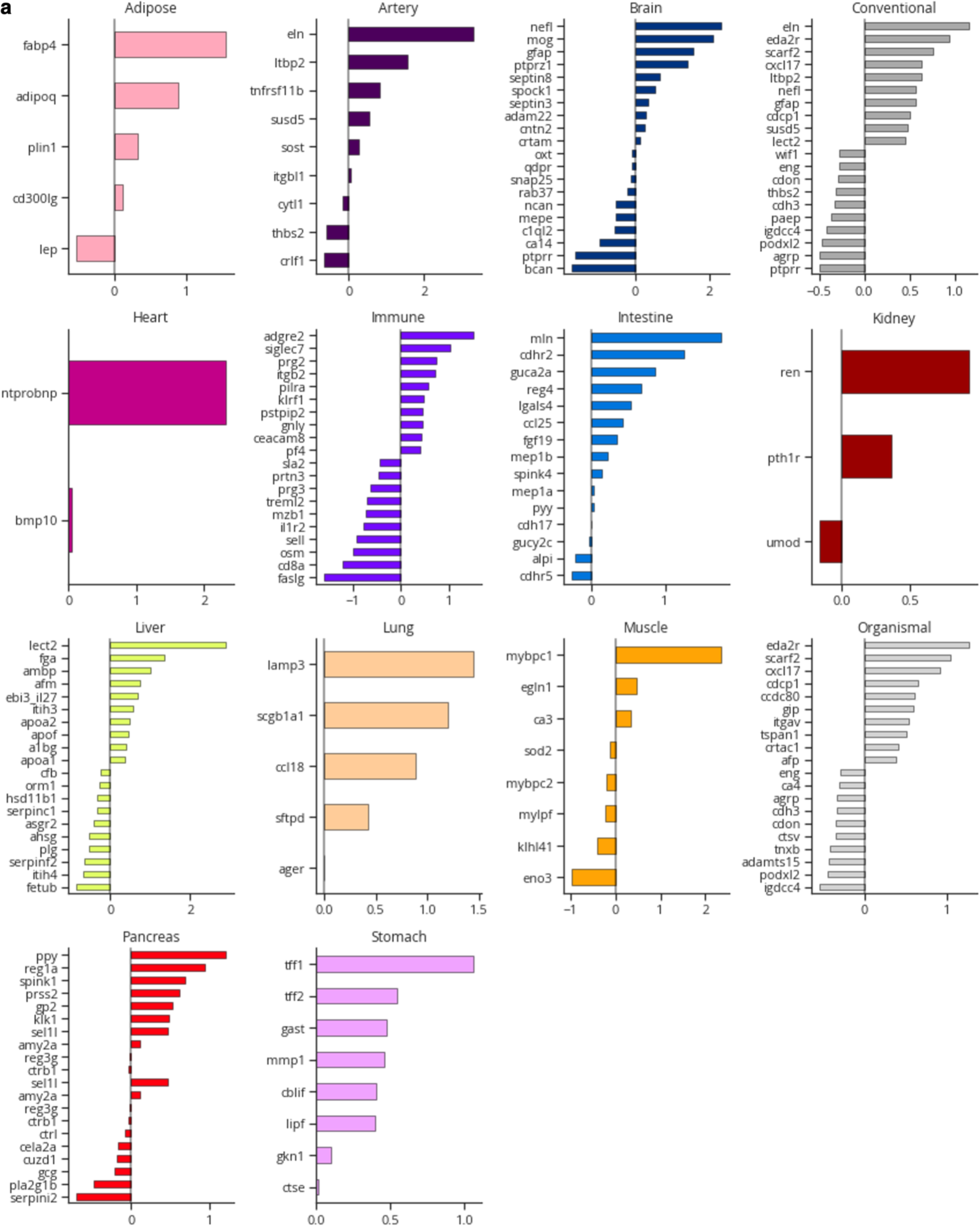
Aging model coefficients. **a,** For all aging models, the top 20 aging model proteins and their weights are shown.

**Extended Data Figure 5.**
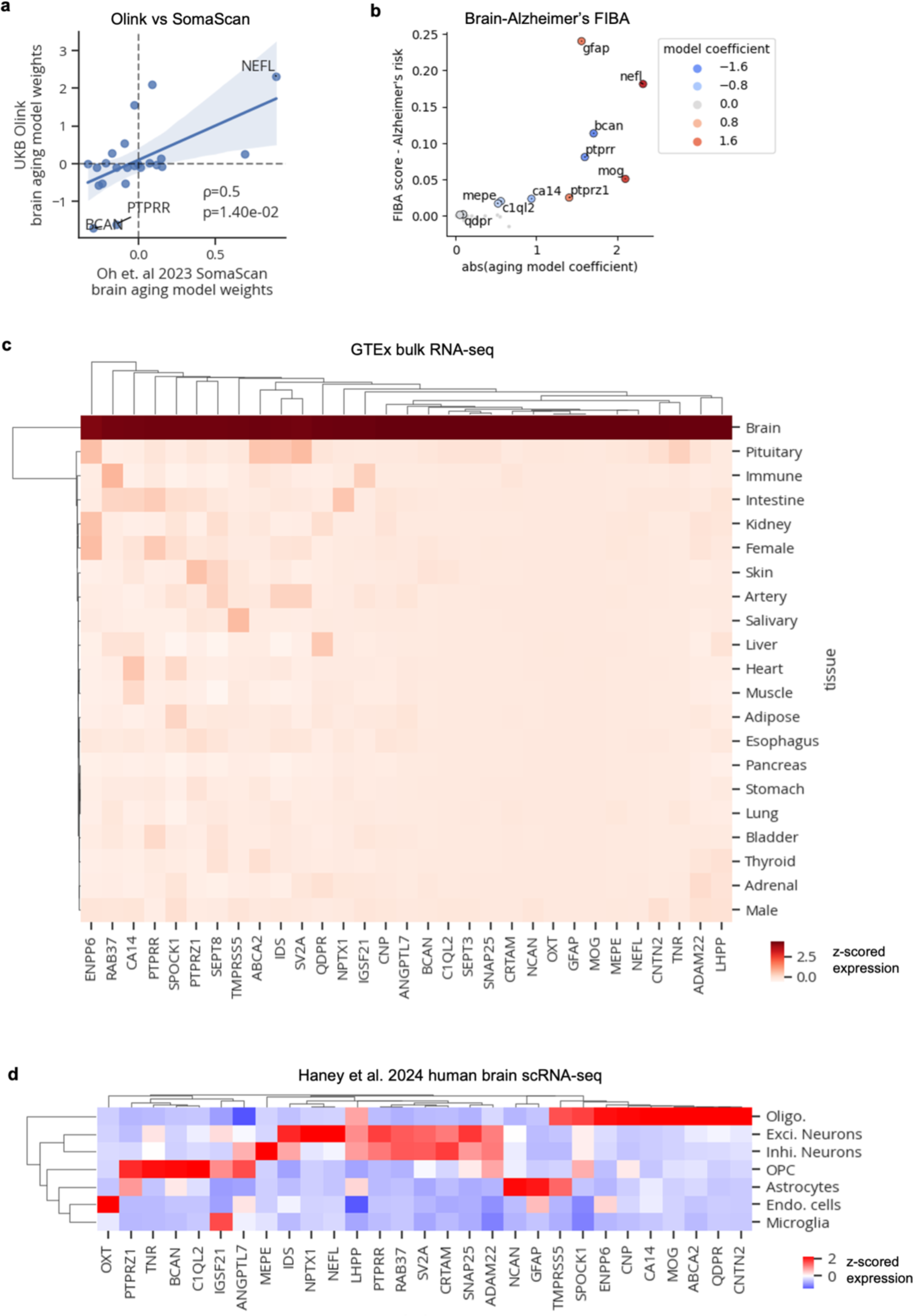
Brain aging proteins. **a,** Scatterplot showing model weights from this study’s Olink-based aging model (y-axis) and Oh et. al 2023’s SomaScan-based brain aging model (x-axis). Spearman correlation and p-value shown. **b,** Scatterplot showing results from feature importance for biological aging (FIBA) algorithm to identify proteins in the brain aging model contributing to the brain age gap’s association with Alzheimer’s disease risk. FIBA score (y-axis) indicates Alzheimer’s disease risk effect size loss after permutation of protein values. X-axis indicates absolute protein weight in the brain aging model. Color indicates protein weight in the brain aging model. **c,** Mean gene expression of brain aging protein-encoding genes in GTEx tissue bulk RNA-seq data. **d,** Mean gene expression of brain aging protein-encoding genes in Haney et al. 2024 human brain scRNA-seq data.

**Extended Data Figure 6.**
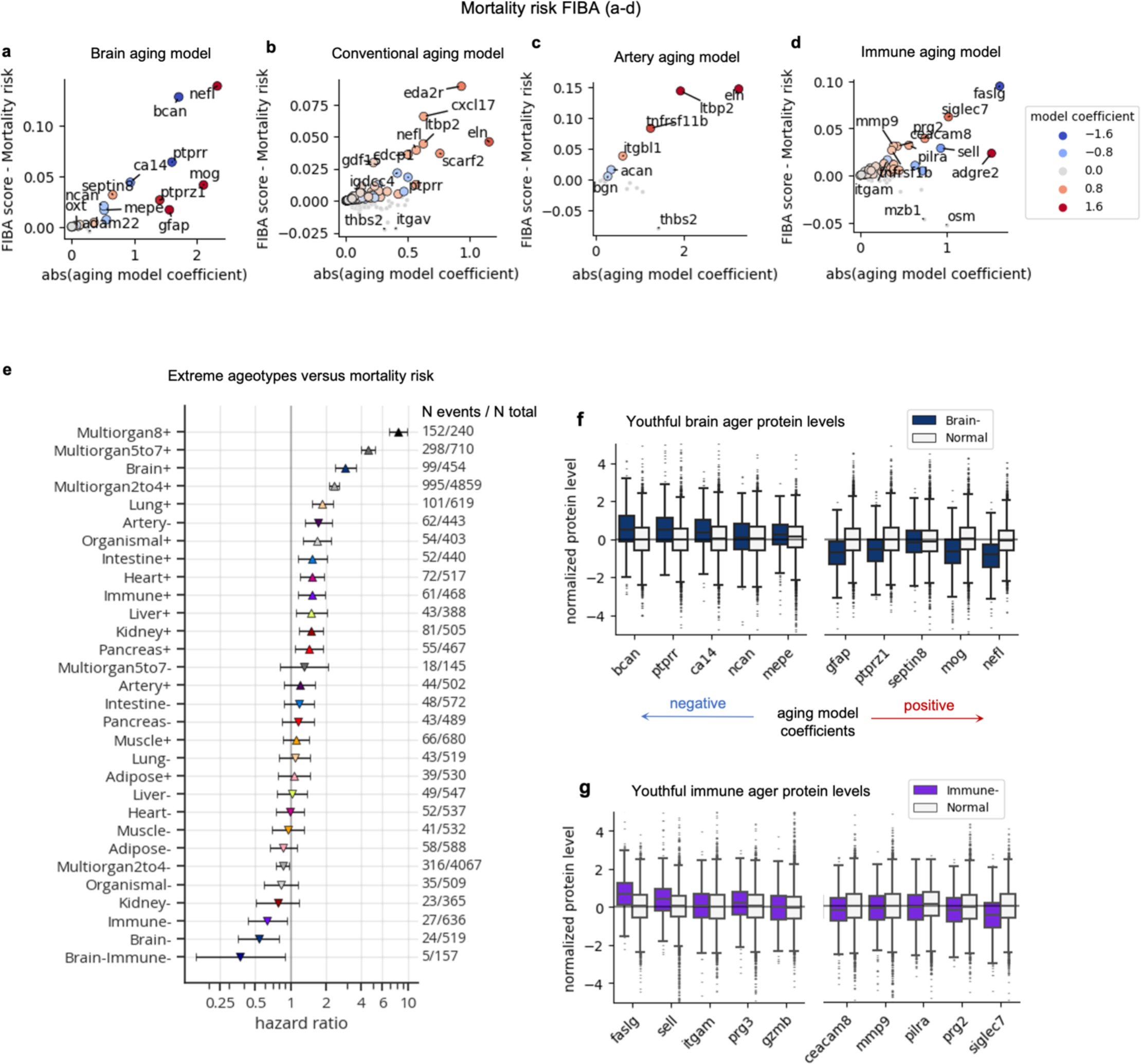
Aging-mortality risk proteins. **a-d,** Scatterplots showing results from feature importance for biological aging (FIBA) algorithm to identify proteins in the brain (**a**), conventional (**b**), artery (**c**), and immune (**d**) aging models that contribute to the model age gap’s association with future mortality risk. FIBA score (y-axis) indicates mortality risk effect size loss after permutation of protein values. X-axis indicates absolute protein weight in the aging model. Color indicates protein weight in the aging model. **e**, Forest plot showing results from Cox proportional hazards regression, testing the associations between extreme ager status and future all-cause mortality risk, controlling for age and sex. Age gap hazard ratios, 95% confidence intervals, number of events out of the total sample size are shown. **f,** Protein levels of youthful brain agers versus normal agers. The top ten (5 decrease with age, 5 increase with age) proteins based on mortality risk FIBA score are shown. Each protein was linearly adjusted for age, sex, and every other protein in the brain aging model before plotting. Proteins are ordered by the aging model coefficient. **g,** As in **f**, but for the immune aging model.

## REFERENCES

1. López-Otín, C., Blasco, M. A., Partridge, L., Serrano, M. & Kroemer, G. Hallmarks of aging: An expanding universe. Cell 186, 243–278 (2023).

2. Ahadi, S. et al. Personal aging markers and ageotypes revealed by deep longitudinal profiling. Nat. Med. 26, 83–90 (2020).

3. Tian, Y. E. et al. Heterogeneous aging across multiple organ systems and prediction of chronic disease and mortality. Nat. Med. 29, 1221–1231 (2023).

4. Oh, H. S.-H. et al. Organ aging signatures in the plasma proteome track health and disease. Nature 624, 164–172 (2023).

5. Raghav Sehgal et al. Systems Age: A single blood methylation test to quantify aging heterogeneity across 11 physiological systems. bioRxiv 2023.07.13.548904 (2023) doi:10.1101/2023.07.13.548904.

6. Herndon, L. A. et al. Stochastic and genetic factors influence tissue-specific decline in ageing C. elegans. Nature 419, 808–814 (2002).

7. Schaum, N. et al. Ageing hallmarks exhibit organ-specific temporal signatures. Nature 583, 596–602 (2020).

8. Rando, T. A. & Wyss-Coray, T. Asynchronous, contagious and digital aging. Nat. Aging 1, 29–35 (2021).

9. Chia-Ling Kuo et al. Proteomic aging clock (PAC) predicts age-related outcomes in middle-aged and older adults. medRxiv 2023.12.19.23300228 (2023) doi:10.1101/2023.12.19.23300228.

10. van Dyck Christopher H., et al. Lecanemab in Early Alzheimer’s Disease. N. Engl. J. Med. 388, 9–21 (2023).

11. Pontecorvo, M. J. et al. Association of Donanemab Treatment With Exploratory Plasma Biomarkers in Early Symptomatic Alzheimer Disease: A Secondary Analysis of the TRAILBLAZER-ALZ Randomized Clinical Trial. JAMA Neurol. 79, 1250–1259 (2022).

12. Miller Timothy M., et al. Trial of Antisense Oligonucleotide Tofersen for SOD1 ALS. N. Engl. J. Med. 387, 1099–1110 (2022).

13. Irwin, K. E., Sheth, U., Wong, P. C. & Gendron, T. F. Fluid biomarkers for amyotrophic lateral sclerosis: a review. Mol. Neurodegener. 19, 9 (2024).

14. Guo, Y. et al. Plasma proteomic profiles predict future dementia in healthy adults. *Nat*. Aging 4, 247–260 (2024).

15. Haney, M. S. et al. APOE4/4 is linked to damaging lipid droplets in Alzheimer’s disease microglia. Nature 628, 154–161 (2024).

16. Fujita, S. et al. Characterization of Brain Volume Changes in Aging Individuals With Normal Cognition Using Serial Magnetic Resonance Imaging. *JAMA Netw*. Open 6, e2318153– e2318153 (2023).

17. Hahn, O. et al. Atlas of the aging mouse brain reveals white matter as vulnerable foci. Cell 186, 4117–4133.e22 (2023).

18. Jonas Morin, Yves Rolland, Heike A. Bischoff-Ferrari, Alejandro Ocampo, & Kevin Perez. Association between prescription drugs and all-cause mortality risk in the UK population. medRxiv 2024.03.08.24303967 (2024) doi:10.1101/2024.03.08.24303967.

19. Faubion, S. S., Kuhle, C. L., Shuster, L. T. & Rocca, W. A. Long-term health consequences of premature or early menopause and considerations for management. Climacteric 18, 483– 491 (2015).

20. Frakes, A. E. et al. Four glial cells regulate ER stress resistance and longevity via neuropeptide signaling in C. elegans. Science 367, 436–440 (2020).

21. Sheng, L. et al. Ensheathing glia promote increased lifespan and healthy brain aging. Aging Cell 22, e13803 (2023).

22. Tokizane, K., Brace, C. S. & Imai, S. DMHPpp1r17 neurons regulate aging and lifespan in mice through hypothalamic-adipose inter-tissue communication. Cell Metab. 36, 377–392.e11 (2024).

23. Koren, T. et al. Insular cortex neurons encode and retrieve specific immune responses. Cell 184, 5902–5915.e17 (2021).

24. Osterhout, J. A. et al. A preoptic neuronal population controls fever and appetite during sickness. Nature 606, 937–944 (2022).

25. Poller, W. C. et al. Brain motor and fear circuits regulate leukocytes during acute stress. Nature 607, 578–584 (2022).

26. Cathomas, F. et al. Circulating myeloid-derived MMP8 in stress susceptibility and depression. Nature 626, 1108–1115 (2024).

27. Jin, H., Li, M., Jeong, E., Castro-Martinez, F. & Zuker, C. S. A body–brain circuit that regulates body inflammatory responses. Nature (2024) doi:10.1038/s41586-024-07469-y.

28. Bonaccio, M. et al. Age-sex–specific ranges of platelet count and all-cause mortality: prospective findings from the MOLI-SANI study. Blood 127, 1614–1616 (2016).

29. Dagfinn Aune et al. BMI and all cause mortality: systematic review and non-linear dose-response meta-analysis of 230 cohort studies with 3.74 million deaths among 30.3 million participants. BMJ 353, i2156 (2016).

30. Karaphillis, E., Goldstein, R., Murphy, S. & Qayyum, R. Serum alanine aminotransferase levels and all-cause mortality. Eur. J. Gastroenterol. Hepatol. 29, (2017).

31. Ulrich-Lai, Y. M. & Herman, J. P. Neural regulation of endocrine and autonomic stress responses. Nat. Rev. Neurosci. 10, 397–409 (2009).

32. Wang, F. et al. Myelin degeneration and diminished myelin renewal contribute to age-related deficits in memory. Nat. Neurosci. 23, 481–486 (2020).

33. Depp, C. et al. Myelin dysfunction drives amyloid-β deposition in models of Alzheimer’s disease. Nature 618, 349–357 (2023).

34. Blanchard, J. W. et al. APOE4 impairs myelination via cholesterol dysregulation in oligodendrocytes. Nature 611, 769–779 (2022).

35. Nott, A. et al. Brain cell type–specific enhancer-promoter interactome maps and disease risk association. Science eaay0793 (2019) doi:10.1126/science.aay0793.

36. Pedregosa, F. et al. Scikit-learn: Machine Learning in Python. J. Mach. Learn. Res. 12, 2825–2830 (2011).

37. Davidson-Pilon, Cameron. (2022). lifelines, survival analysis in Python (v0.27.0). Zenodo. 10.5281/zenodo.6359609.

38. Seabold, S. & Perktold, J. Statsmodels: Econometric and Statistical Modeling with Python. in *Proceedings of the 9th Python in Science Conference* (eds. Walt, S. van der & Millman, J.) 92–96 (2010). doi:10.25080/Majora-92bf1922-011.

